# SMuRF: Portable and accurate ensemble-based somatic variant calling

**DOI:** 10.1101/270413

**Authors:** Weitai Huang, Yu Amanda Guo, Karthik Muthukumar, Probhonjon Baruah, Meimei Chang, Anders Jacobsen Skanderup

## Abstract

**Summary:** SMuRF is an ensemble method for prediction of somatic point mutations (SNVs) and small insertions/deletions (indels) in cancer genomes. The method integrates predictions and auxiliary features from different somatic mutation callers using a Random Forest machine learning approach. SMuRF is trained on community-curated tumor whole genome sequencing data, is robust across cancer types, and achieves improved accuracy for both SNV and indel predictions of genome and exome-level data. The software is user-friendly and portable by design, operating as an add-on to the community-developed bcbio-nextgen somatic variant calling pipeline.

## INTRODUCTION

Identification of somatic mutations from matched tumor and normal samples is challenged by sequencing and alignment artefacts as well as the heterogeneous composition and mutational processes of tumors. Recent studies have revealed low concordance between existing methods for somatic variant calling (1-4). Additionally, a benchmark study demonstrated that the accuracy of a given somatic mutation calling algorithm can vary extensively across different workflows and pipelines (5). Parameters influencing this variation may be choice of alignment algorithm, use of local re-alignment, as well as configuration of a multitude of post-processing filters. Furthermore, somatic mutation calling algorithms are often trained and evaluated on simulated tumor data and/or whole exome sequencing (WES) data, and it is less clear how well they perform in a genome-wide setting. Indeed, a recent benchmark study demonstrated that many methods and pipelines have low accuracy when applied to tumor whole-genome sequencing data (5).

Previous studies have used the consensus of multiple callers to improve the accuracy of somatic variant calling (6,7). Taking this one step further, a machine learning based ensemble method may combine multiple mutation callers with auxiliary sequence and alignment features to improve mutation calling accuracy (8). While such approaches may improve accuracy, they are generally not portable. The end-user must identify and obtain a suitable training and testing datasets, need to have knowledge of machine learning, and must re-fit the model whenever parameters of the workflow changes (Fig. 1A). There is therefore a need for accurate ensemble approaches for somatic mutation calling that can be ported between research groups.

**Figure 1:**
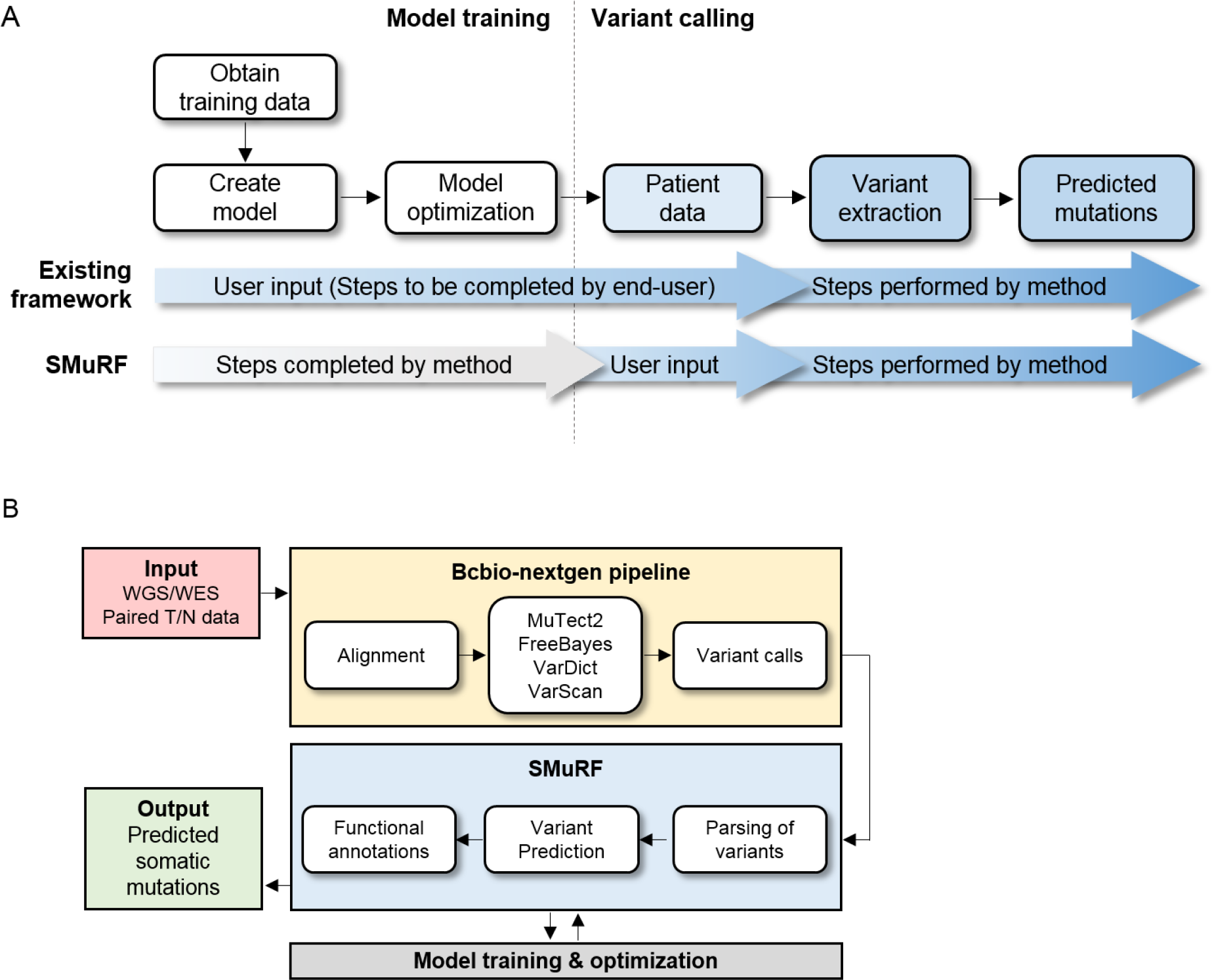
Overview of approach. (A) General steps involved in an ensemble method for somatic mutation calling. In existing frameworks, end-users must obtain data to train the model before variant calling can be performed. In SMuRF, the model has already been pre-trained and end-users just need to provide variant calls from the bcbio-nextgen workflow. (B) Schematic workflow of somatic mutation analysis with SMuRF. Users provide input data of .fastq files for the bcbio-nextgen somatic variant calling pipeline. Variant call files are used in the SMuRF ensemble somatic mutation calling step. Output is the set of predicted somatic SNVs and indels together with their SMuRF confidence scores.

Bcbio-nextgen is a highly popular (with almost 1000 code commits from over 50 contributors in 2017) community developed next generation sequencing analysis pipeline that is fully automated, validated, and scalable (9). The default bcbio-nextgen somatic variant calling pipeline employs four mutation callers: MuTect2 (10), Freebayes (11), VarDict (12) and VarScan (13), and offers a simple consensus vote to select the final set of somatic mutation variant calls. Here we developed a **S**omatic **Mu**tation calling method using a **R**andom **F**orest model (SMuRF), which combines the predictions from these individual mutation callers with auxiliary alignment and mutation features using supervised machine learning (Fig. 1B). By integrating SMuRF directly with the bcbio-nextgen framework, SMuRF is portable by design and require minimal efforts to setup and run. SMuRF is trained on a gold standard set of mutation calls that has been curated by the International Cancer Genome Consortium (ICGC) community using deep (>100x) whole genome sequencing (WGS) of two tumors (5). We show that SMuRF is highly accurate and outperforms individual somatic mutation callers as well as simple consensus voting. We demonstrate that the model is robust across different cancer types and is highly accurate for both WGS and WES data.

## MATERIALS AND METHODS

### Input and output

SMuRF requires input files in variant call format (VCF) from the somatic mutation callers MuTect2, Freebayes, VarDict, and VarScan as generated by the default ‘variant2’ bcbio-nextgen pipeline (9). The output from a SMuRF analysis is a set of data tables containing predicted SNVs and indels together with their associated confidence scores. The data tables can easily be converted into the mutation annotation format (MAF) which is compatible with other programs for downstream analysis.

### Implementation

SMuRF is written in R and is freely available as an R package. First, VCF files from bcbio-nextgen are processed to extract variant features which include base and mapping quality scores and other features provided by each of the four callers. SMuRF then uses a Random Forest (h2o R package) classifier to accurately predict SNVs and indels given these features. Feature extraction and prediction of somatic variants takes ~15 minutes for tumor-normal WGS data on a standard computer (4 CPUs, 16GB RAM). To ensure future compatibility between SMuRF and bcbio-nextgen, the performance of SMuRF can be re-assesed, and model potentially re-trained, with future bcbio-nextgen releases. Information about compability of SMuRF and bcio-nextgen versions will be documented on the SMuRF Github page.

### Annotation of somatic mutations

The “cdsannotation” function in the SMuRF package adds annotations for predicted mutations in the coding regions by merging annotation provided by the bcbio-nextgen pipeline (by SnpEff) (14).

### Training datasets

The Random Forest model was trained on matched tumor-normal WGS data from a chronic lymphocytic leukemia (CLL) patient and a medulloblastoma (MB) patient, where the true somatic mutations have been identified through an extensive community curation procedure by the International Cancer Genome Consortium (ICGC) (5). For model training, we down-sampled both tumor datasets to sequence coverages typically generated in WGS studies (CLL: tumor/normal ~40x/~50x; MB: ~75x/~50x). Both tumors had a high tumor purity (cancer-cell fraction; ~0.92 for CLL and ~0.98 for MB).

### Feature selection, optimization and model training

In the selection of features, potentially predictive alignment and variant features were extracted from the vcf-files. Features directly dependent on sequencing coverage, such as allele counts and read depth, were excluded. The features were evaluated by the random forest algorithm and ranked based on their relative importance. The most consistent features (with a geometric mean ranking <10) when training on the CLL or MB tumor alone were selected. Overall, 9 and 10 features were selected for the SNV (Table 1) and indel (Table 2) models, respectively. The maximum number of randomly selected features (*mtries*) and maximum tree depth (*depth*) in the Random Forest model were determined by a grid search strategy using 5-fold cross validation (SNV: *mtry* = 5, *depth* = 11; indel: *mtry* = 4, *depth* = 10).

**Table 1:** Summary of SNV features. Summary of the selected features for the SMuRF SNV model. Features were obtained from the corresponding callers. The percent importance indicates the relative importance of the particular feature in the overall prediction model and rank indicates the overall ranking of the feature. The percent importance and ranks were derived for the CLL and MB dataset independently. Mean ranks are calculated based on the geometric mean (G.mean) between the ranks of the CLL and MB dataset.

| No. | Caller | Features | %importance |  | Ranks |  | G.mean<sup>#</sup> | Description |
| --- | --- | --- | --- | --- | --- | --- | --- | --- |
|  |  |  | CLL | MB | CLL | MB |  |  |
| 1 | MuTect2 | MQ | 0.14 | 0.19 | 2 | 1 | 1.41 | RMS Mapping Quality |
| 2 | MuTect2 | TLOD | 0.22 | 0.13 | 1 | 2 | 1.41 | Tumor LOD score |
| 3 | FreeBayes | MQMR | 0.10 | 0.08 | 3 | 5 | 3.87 | Mean mapping quality of observed reference alleles |
| 4 | MuTect2 | MQRankSum | 0.06 | 0.12 | 7 | 3 | 4.58 | Z-score From Wilcoxon rank sum test of Alt vs. Ref read mapping qualities |
| 5 | VarDict | SSF | 0.09 | 0.07 | 4 | 6 | 4.90 | p-value |
| 6 | FreeBayes | MQM | 0.05 | 0.12 | 8 | 4 | 5.66 | Mean mapping quality of observed alternate alleles |
| 7 | VarScan | SSC | 0.08 | 0.06 | 5 | 8 | 6.32 | Somatic score in Phred scale (0-255) derived from somatic p-value |
| 8 | MuTect2 | NLOD | 0.04 | 0.07 | 11 | 7 | 8.77 | Normal LOD score |
| 9 | VarScan | SPV | 0.07 | 0.01 | 6 | 15 | 9.49 | Fisher's Exact Test P-value of tumor versus normal for Somatic/LOH calls |

<sup>#</sup>Geometric mean calculated between CLL and MB ranks

**Table 2:** Summary of indel features. Summary of the selected features for the SMuRF INDEL model. Features were obtained from the corresponding callers. The percent importance indicates the relative importance of the particular feature in the overall prediction model and rank indicates the overall ranking of the feature. The percent importance and ranks were derived for the CLL and MB dataset independently. Mean ranks are calculated based on the geometric mean (G.mean) between the ranks of the CLL and MB dataset.

| No. | Caller | Features | %importance |  | Ranks |  | G.mean<sup>#</sup> | Description |
| --- | --- | --- | --- | --- | --- | --- | --- | --- |
|  |  |  | CLL | MB | CLL | MB |  |  |
| 1 | VarScan | SPV | 0.23 | 0.15 | 1 | 3 | 1.73 | Fisher's Exact Test P-value of tumor versus normal for Somatic/LOH calls |
| 2 | VarScan | SSC | 0.08 | 0.17 | 4 | 1 | 2.00 | Somatic score in Phred scale (0-255) derived from somatic p-value |
| 3 | MuTect2 | TLOD | 0.09 | 0.08 | 2 | 6 | 3.46 | Tumor LOD score |
| 4 | MuTect2 | NLOD | 0.06 | 0.15 | 8 | 2 | 4.00 | Normal LOD score |
| 5 | VarDict | SSF | 0.08 | 0.06 | 3 | 7 | 4.58 | p-value |
| 6 | MuTect2 | MQ | 0.06 | 0.09 | 6 | 4 | 4.90 | RMS Mapping Quality |
| 7 | VarDict | MSI | 0.07 | 0.08 | 5 | 5 | 5.00 | Micro-satellite |
| 8 | FreeBayes | LEN | 0.06 | 0.06 | 7 | 8 | 7.48 | Allele Length |
| 9 | MuTect2 | MQRankSum | 0.05 | 0.05 | 10 | 9 | 9.49 | Z-score From Wilcoxon rank sum test of Alt vs. Ref read mapping qualities |
| 10 | VarDict | SOR | 0.05 | 0.03 | 9 | 10 | 9.49 | Odds ratio |

<sup>#</sup>Geometric mean calculated between CLL and MB ranks

### Evaluation of model accuracy

We trained our model on 70% of the combined CLL and MB data using five-fold crossvalidation, withholding the remaining 30% as independent test data. The four individual variant callers were run with default parameters in the bcbio-nextgen pipeline. The accuracy of the model and individual mutation callers were evaluated with the F1-score, which is the harmonic mean between the precision and recall 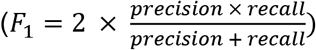, where 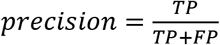 and 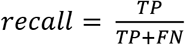. Precision-recall curves were plotted to visualize the performance of SMuRF compared to the individual callers. We plotted precision-recall curves using the confidence score provided by each method: SMuRF confidence score, MuTect2 tumor log-odds score, Freebayes log-odds score, VarDict SSF score, and VarScan SSC score. The performance of mutation calls identified by the majority-vote (intersection of at least 1, 2, 3 or all 4 callers) were also evaluated.

### Generation of tumor samples with lower purity levels

We generated tumor samples with lower purity by adding normal reads from the corresponding matched normal bam files into the tumor bam files. This provided lowered purity at two settings: tumor + 20x normal reads (20x contamination), and tumor + 40x normal reads (40x contamination). We also further down-sampled the normal samples to 30x sequence coverage (unbalanced) to simulate samples with unbalanced tumor/normal sequence coverage.

### Comparison of concordance for WGS and WES data

We analyzed an in-house colorectal cancer as well as The Cancer Genome Altas (TCGA) liver cancer cohort. 10 patient samples were randomly selected to evaluate the concordance of the mutation calls between WGS and WES platforms. The WGS/WES concordance was calculated based on the Jaccard index, 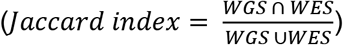, which is the ratio of the intersection over the union of the WGS and WES calls in the coding regions.

## RESULTS AND DISCUSSION

### SMuRF is accurate for SNV and indel calling

SMuRF uses separate models for SNV and indel prediction. The models incorporate the top predictive features extracted from the individual mutation callers (Table 1 and 2). To benchmark the performance of SMuRF in comparison with individual variant calling methods, we evaluated the accuracy (precision and recall) on an independent test set. The accuracy recorded for individual mutation callers in our comparison was comparable to the recent benchmark by Alioto et al. using the same dataset (5). While the best method in this benchmark obtained an F1-score of 0.79 and 0.65 for SNVs and indels respectively, SMuRF achieved an F1-score of 0.91 and 0.77 (using independent test data). The accuracy of SMuRF was noticeably higher than each of the four individual mutation callers as well as simple majority voting models (Fig. 2A-B). While all of the individual callers could recover most of the true SNVs (>90% recall), this came at the cost of very low precision (<20%). In contrast, SMuRF could recover 90% of the true SNVs at 92% precision and recover 77% of the true indels at 76% precision.

**Figure 2:**
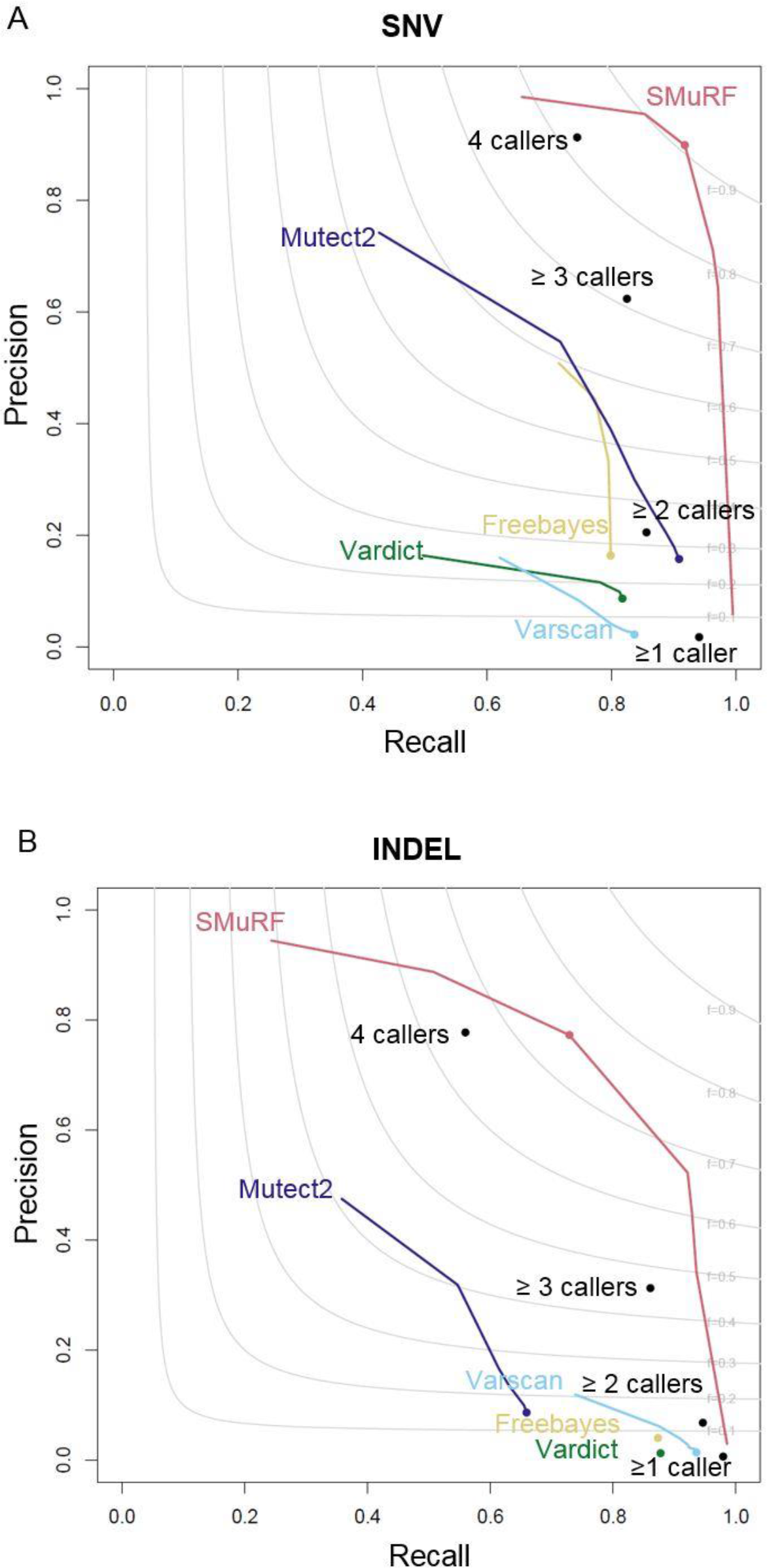
Performance of SMuRF. Precision-recall profiles for individual somatic mutation callers and SMuRF evaluated on (A) SNV and (B) indels. SMuRF performance was evaluated from independent test data. Curves show the performance of the individual algorithms under different variant score thresholds (MuTect2 tumor log-odds score, Freebayes log-odds score, VarDict SSF score, VarScan SSC score, and SMuRF confidence score). Solid points refer to the default performance of the caller in the bcbio-nextgen workflow. The black solid points denote the accuracy of concordant calls identified by the majority-voting scheme in bcbio-nextgen (at least 1, 2, 3, or 4 callers). The grey curves show F1 scores as a function of recall and precision.

In order to ensure that SMuRF is robust to variations in sequencing coverage and tumor purity, we performed spike-in of normal reads into the two tumor samples (contamination of 20x and 40x normal reads). We also down-sampled the sequencing coverage of the normal sample to 30x (“unbalanced”) so as to simulate unbalanced tumor/normal genome sequencing (Fig. 3A).

**Figure 3:**
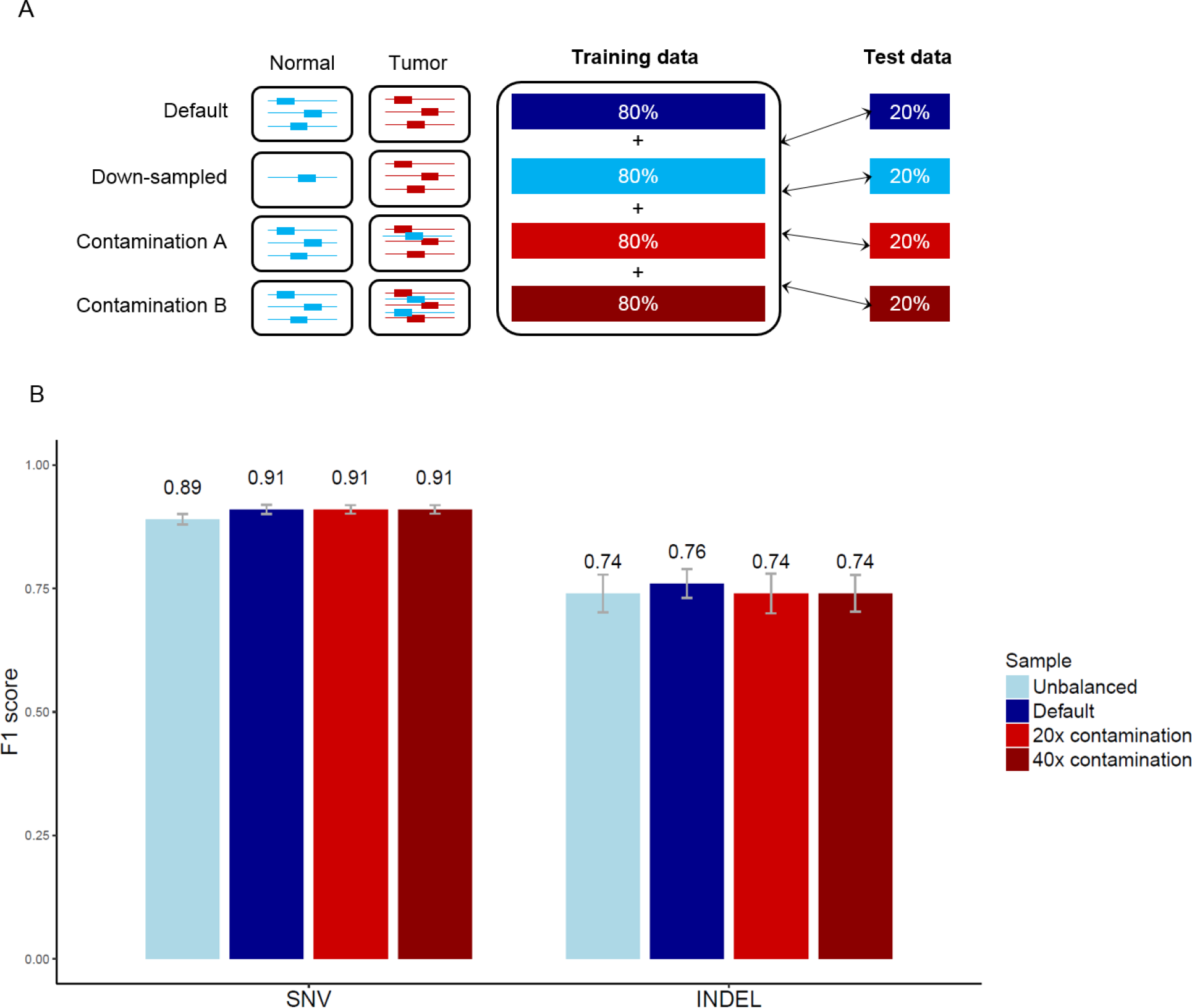
Robustness of SMuRF. (A) Different conditions were generated from the datasets: original data (Default), normal sample down-sampled to 30x coverage (Unbalanced), lowered purity tumor samples spiked in with either 20x or 40x normal reads respectively (20x & 40x contamination). (B) F1 accuracy scores under each condition when evaluated on independent test data (20%). Scores were obtained from 5-fold cross-validation repeated 10 times.

We trained SMuRF on the union of these datasets using 5-fold cross-validation, withholding 20% of positions as independent test data. Neither lower tumor sample purity nor low sequencing coverage severely affected the accuracy of SMuRF (SNV: F1=0.91 for default, F1=0.89 for unbalanced, F1=0.91 for 20x contamination, F1=0.91 for 40x contamination; indels: F1=0.76 for default, F1=0.74 for unbalanced, F1=0.74 for 20x contamination, F1=0.74 for 40x contamination). These results support that SMuRF is robust to variations in sequencing coverage and tumor purity.

### SMuRF is robust across different cancer types

A critical feature of a somatic variant caller is its ability to accurately predict somatic mutations across different cancer types, which may have unique mutational signatures and processes. Hence, we evaluated the extent that SMuRF predictions can generalize across cancer types. We therefore trained SMuRF on the CLL data and tested it on the MBL data, and vice versa (Fig. 4A). Cross-cancer evaluation between the CLL and MB datasets showed minimal changes in performance for SNV (F1: 0.87 vs. 0.86) and slight variation for indel (F1: 0.57 vs. 0.64) prediction respectively (Fig. 4B). This indicates that SMuRF is not over-fitted to a particular cancer type and may be applicable for somatic mutation predictions in other cancer types.

**Figure 4:**
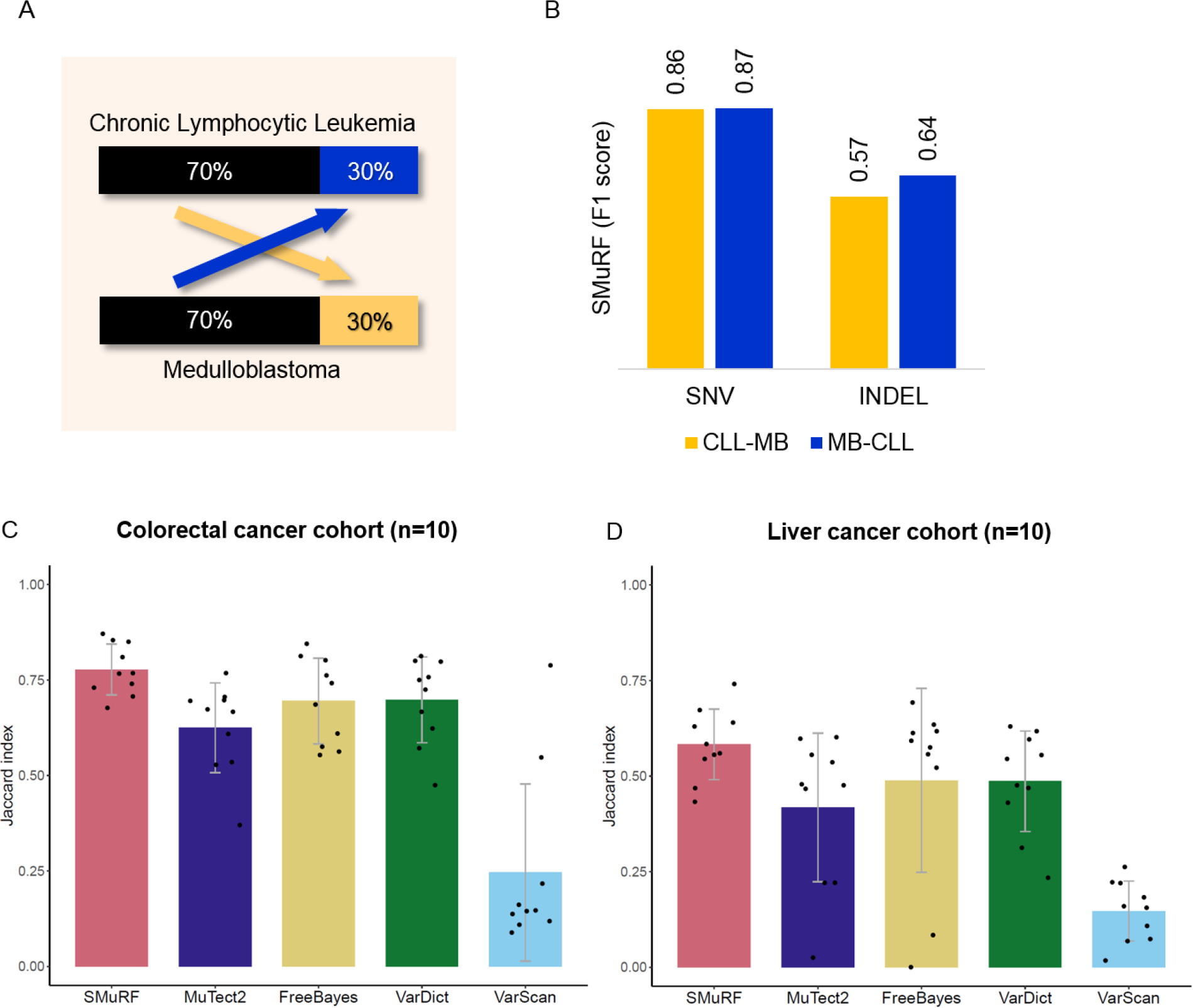
Performance across cancer types. Training and testing on datasets from different cancer types. (A) Schematic of training and testing on the CLL and MB datasets and vice versa using 70-30 (train-test) as compared to the respective SMuRF (SNV or Indel) model and the performance evaluated by the F1 score (B). (C & D) Concordance of SNV coding mutations between WGS and WES samples. 10 Colorectal Cancer (C) and Liver Cancer (D) patients were randomly chosen. The barplots shows the average Jaccard index for WES compared to WGS variants in the coding region for each of the callers. Error bars indicate the standard deviation of the mean and the points indicate the Jaccard index achieved by each sample. The number of calls analyzed in each caller were fixed to the total number of calls in SMuRF and selected based on the top-K mutations ranked by the respective caller features: MuTect2 tumor log-odds score, Freebayes log-odds score, VarDict SSF score & VarScan SSC score.

Secondly, we analyzed the extent that SMuRF predicts the same somatic mutations in tumor samples profiled with both WES and WGS. We first selected 10 random in-house sequenced colorectal cancer samples with both WES and WGS data. Firstly, SMuRF was able to correctly predict mutations at a genome-wide rate (~10k-150k) expected from previous studies of colorectal cancer (~3k-300k per genome (15)). Secondly, we compared the overlap between WES and WGS variant calls, restricting analysis to variants in coding regions (defined by Ensembl). SMuRF achieved an average Jaccard Index concordance of 0.78 between the WES and WGS platforms (Fig. 4C). In order to compare individual callers with SMuRF, we used the top-K predictions from individual callers, where K was determined by the number of variants identified by SMuRF for a given sample. MuTect2, Freebayes, Vardict and VarScan achieved an average concordance of 0.62, 0.69, 0.70 and 0.25 respectively. Similarly, we evaluated 10 randomly selected samples from the public TCGA liver cancer dataset having both WES and WGS data. Here SMuRF predicted ~5k-25k SNVs per tumor genome, which is consistent with the expected mutation rate for liver cancer (~3k-30k per genome (15)). SMuRF achieved a concordance of 0.58 between WES and WGS calls, while the individual callers MuTect2, Freebayes, Vardict and VarScan achieved a concordance of 0.44, 0.49, 0.49 and 0.15 respectively (Fig. 4D).

Overall, SMuRF predicted SNVs at a higher concordance as compared to the individual callers in both datasets. We also observed a similar trend for the indels, with SMuRF achieving the highest concordance (Suppl. Fig. 1). We observed an overall reduction in concordance in the liver cancer dataset compared to the colorectal cancer dataset for both SMuRF and individual callers (Fig. 4C-D & Suppl. Fig. 1). The difference in the concordance of the two cancer datasets may be due to lower sequencing coverage in the liver cancer cohort (100-400x for WES, 40-60x for WGS) as compared to the colorectal cancer cohort (450-600x WES, 60-80x WGS) (Suppl. Fig. 2). The evaluation showed that SMuRF has the ability to analyze not only WGS data but also WES data and is capable of identifying a concordant set of mutations for downstream analyses.

## CONCLUSION

SMuRF identifies somatic mutations with high accuracy by combining the predictions from individual somatic mutation callers with auxiliary alignment and mutation features using supervised machine learning. SMuRF is trained on a gold standard set of mutation calls curated by the scientific community using deep WGS of two tumors (5). We show that SMuRF can predict SNVs and indels with improved accuracy in both WGS and WES data. Furthermore, SMuRF is robust to variation in tumor purity and tumor-normal sequence coverage bias, and can predict mutations with improved accuracy across different cancer types.

By integrating SMuRF directly with the community validated and highly scalable bcbio-nextgen framework, SMuRF is portable by design and require minimal efforts to setup and run. The tight integration with the bcbio-nextgen framework may present a barrier for some users. However, this design ensures that end-users do not need to obtain their own training data and train their own machine learning models, a process that is both time consuming and require knowledge of machine learning. In summary, SMuRF is an accurate, portable, and user-friendly somatic mutation caller, which should benefit both large-scale cancer genomics studies as well as clinical applications.

## AVAILABILITY

SMuRF is written in R and is freely available on GitHub: https://github.com/skandlab/SMuRF. Documentation for SMuRF is available at https://github.com/skandlab/SMuRF/wiki.

## ACKNOWLEDGEMENTS

We thank members of the Skanderup lab (https://www.skandlab.org/) for their support and discussion during the development and pre-release of SMuRF.

## Supplementary Information

**Supplementary figure 1.**
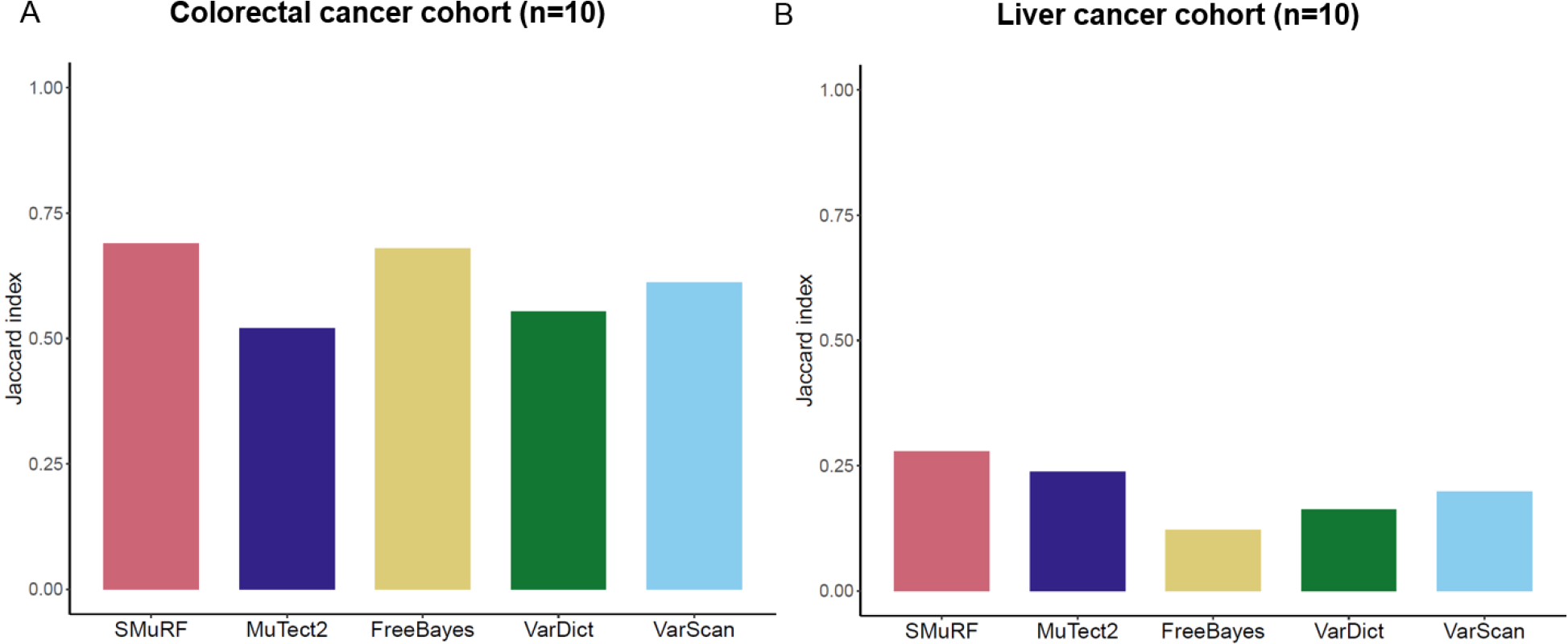
Concordance of INDEL coding mutations between WGS and WXS samples from 10 randomly chosen (A) Colorectal Cancer and (B) Liver Cancer patients. Indels from all 10 samples were pooled together for analysis. The plots shows the Jaccard index for WXS compared to WGS variants in the coding region for each of the callers. The number of calls analyzed in each caller were fixed to the total number of calls in SMuRF and selected based on the top-K mutations ranked by the respective caller features: MuTect2 tumor log-odds score, Freebayes allele length, VarDict SSF score & VarScan SSC score.

**Supplementary figure 2.**
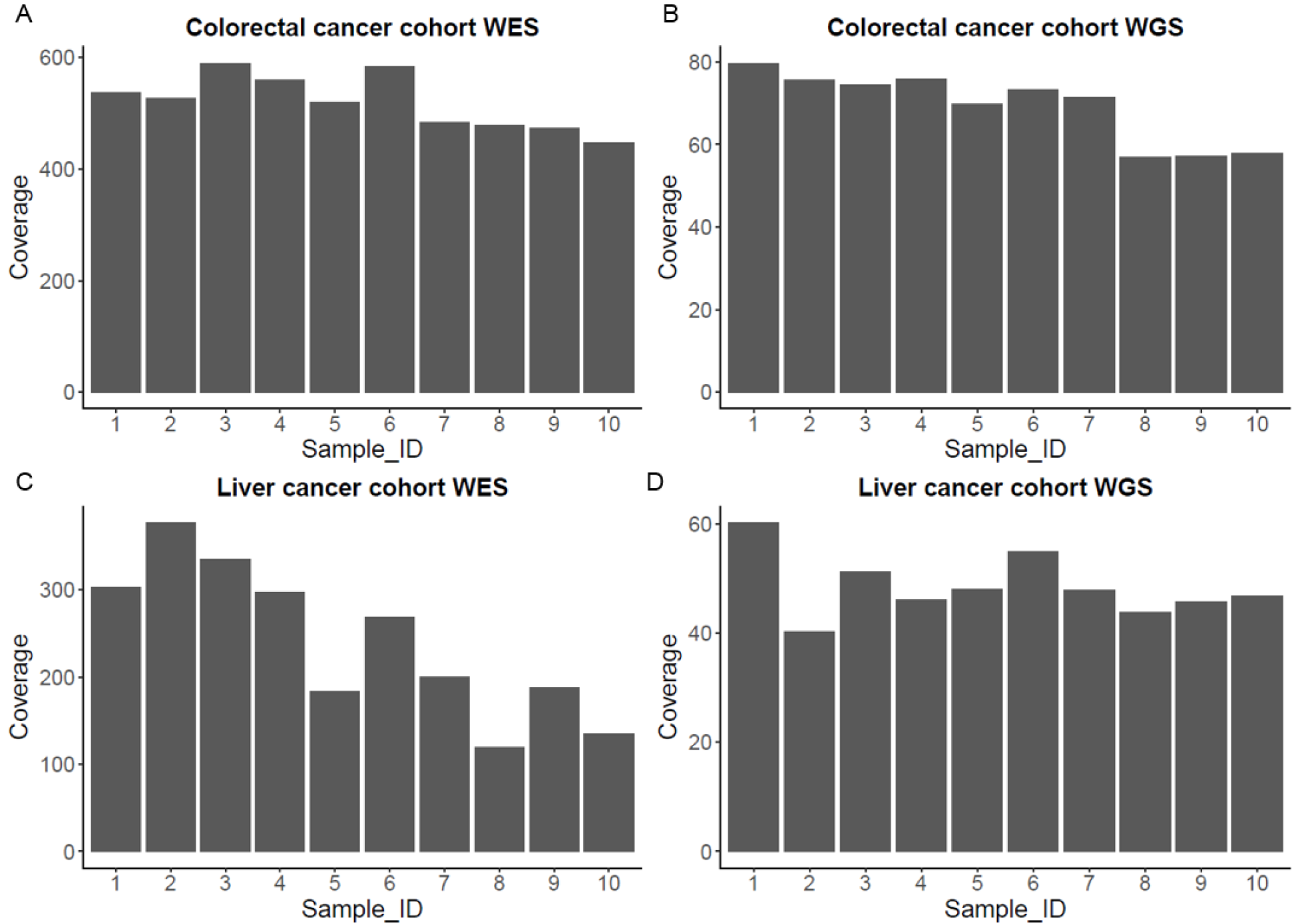
Coverage estimates for WXS and WGS samples from 10 randomly chosen (A & B) Colorectal Cancer and (C & D) Liver Cancer patients respectively. The coverage estimates are calculated from the average of the total read depth from variants in the coding regions from each of the paired normal and tumor samples.

